# Spatial transcriptomics analysis of neoadjuvant cabozantinib and nivolumab in advanced hepatocellular carcinoma identifies independent mechanisms of resistance and recurrence

**DOI:** 10.1101/2023.01.10.523481

**Authors:** Shuming Zhang, Long Yuan, Ludmila Danilova, Guanglan Mo, Qingfeng Zhu, Atul Deshpande, Alexander T.F. Bell, Jennifer Elisseeff, Aleksander S. Popel, Robert A. Anders, Elizabeth M. Jaffee, Mark Yarchoan, Elana J. Fertig, Luciane T. Kagohara

**Affiliations:** Department of Biomedical Engineering, Johns Hopkins University School of Medicine, Baltimore, MD, USA; Department of Immunology, Johns Hopkins University School of Medicine, Baltimore, MD, USA; Bloomberg-Kimmel Immunotherapy Institute, Johns Hopkins University School of Medicine, Baltimore, MD, USA; Department of Oncology, Sidney Kimmel Comprehensive Cancer Center, Johns Hopkins University School of Medicine, Baltimore, MD, USA; Convergence Institute, Johns Hopkins University, Baltimore, MD, USA; Department of Pathology, Johns Hopkins University School of Medicine, Baltimore, MD, USA; Department of Ophthalmology, Johns Hopkins University School of Medicine, Baltimore, MD, USA; Department of Orthopedic Surgery, Johns Hopkins University School of Medicine, Baltimore, MD, USA; Department of Applied Mathematics and Statistics, Johns Hopkins University School of Medicine, Baltimore, MD, USA

**Author notes:** Corresponding authors: Luciane T. Kagohara, 1650 Orleans St – Rm 485, Baltimore, MD 21287, Elana J. Fertig, 550 N Broadway – Rm 1101, Baltimore, MD 21287.

**Keywords:** hepatocellular carcinoma, spatial transcriptomics, therapeutic resistance, neoadjuvant therapy, immunotherapy, tumor recurrence

## Abstract

Novel immunotherapy combination therapies have improved outcomes for patients with hepatocellular carcinoma (HCC), but responses are limited to a subset of patients and recurrence can also occur. Little is known about the inter- and intra-tumor heterogeneity in cellular signaling networks within the HCC tumor microenvironment (TME) that underlie responses to modern systemic therapy. We applied spatial transcriptomics (ST) profiling to characterize the tumor microenvironment in HCC resection specimens from a clinical trial of neoadjuvant cabozantinib, a multi-tyrosine kinase inhibitor that primarily blocks VEGF, and nivolumab, a PD-1 inhibitor in which 5 out of 15 patients were found to have a pathologic response. ST profiling demonstrated that the TME of responding tumors was enriched for immune cells and cancer associated fibroblasts (CAF) with pro-inflammatory signaling relative to the non-responders. The enriched cancer-immune interactions in responding tumors are characterized by activation of the PAX5 module, a known regulator of B cell maturation, which colocalized with spots with increased B cell markers expression suggesting strong activity of these cells. Cancer-CAF interactions were also enriched in the responding tumors and were associated with extracellular matrix (ECM) remodeling as there was high activation of FOS and JUN in CAFs adjacent to tumor. The ECM remodeling is consistent with proliferative fibrosis in association with immune-mediated tumor regression. Among the patients with major pathologic response, a single patient experienced early HCC recurrence. ST analysis of this clinical outlier demonstrated marked tumor heterogeneity, with a distinctive immune-poor tumor region that resembles the non-responding TME across patients and was characterized by cancer-CAF interactions and expression of cancer stem cell markers, potentially mediating early tumor immune escape and recurrence in this patient. These data show that responses to modern systemic therapy in HCC are associated with distinctive molecular and cellular landscapes and provide new targets to enhance and prolong responses to systemic therapy in HCC.

## BACKGROUND

Hepatocellular carcinoma (HCC) is one of the most common causes of cancer associated deaths globally and the most rapidly rising cause of cancer death in the United States [1–3]. Immune checkpoint inhibitors (ICIs) that target programmed cell death protein-1 (PD1) have modest clinical activity as monotherapy in HCC but may be more effective in combination with other therapeutic agents, including anti-angiogenic therapies. The combination of bevacizumab (an anti-VEGF antibody) plus atezolizumab (an ICI targeting the PD1 axis) was recently established as a preferred first line standard of care for patients with unresectable HCC and clinical efficacy has also been reported with multiple other anti-angiogenic/immune checkpoint inhibitor (ICI) combinations (cabozantinib + atezolizumab, and apatinib + camrelizumab) [4–10]. While anti-angiogenic plus ICI combinations have demonstrated clinical benefit, a significant proportion of patients does not respond to such therapies, and biomarkers to predict response and determine which patients will truly benefit from the treatment are currently not available [11, 12].

Window of opportunity studies provide an unparalleled opportunity to elucidate mechanisms of response and resistance to systemic therapy, yielding abundant tissue for deep interrogation of the tumor microenvironment (TME) in patients receiving neoadjuvant systemic therapy. We recently reported the results of a window of opportunity clinical trial of angiogenic/ICI therapy in which HCC patients were treated with 8 weeks of cabozantinib and nivolumab (CABO/NIVO) followed by attempted surgical resection (NCT03299946) [10]. In this clinical study, 5 out of 15 patients achieved pathologic responses, with outstanding disease-free survival noted among patients with a pathologic response. Spatial proteomics profiling of the HCC surgical specimens identified an enriched immune effector infiltrate, and reduced immunosuppressive macrophages, among patients with pathologic response to CABO/NIVO [10]. Nevertheless, molecular pathways that underlie these immune interactions as well as the tumor intrinsic mechanisms of resistance to ICIs in HCC are not clear. In this current study, we performed a high-dimensional, unbiased profiling of patients enrolled in our window of opportunity clinical trial of cabozantinib and nivolumab in HCC to uncover the intrinsic molecular and cellular tumor features of the tumor microenvironment that underlie differential responses to systemic therapy.

The recent development of technologies that provide spatially resolved gene expression data introduced powerful methods to profile the TME and understand how tumor intrinsic features are associated with the distribution of other crucial cell types for tumor development and response to therapies [13]. Using spatial transcriptomics (ST), it is possible to examine samples molecular and cellular compositions, and interactions among the different cellular components as it maintains tissue architecture [14]. Thus, ST provides spatial visualization of immune cells distribution in relation to the cancer cells and the correlation with the molecular profile that drive the infiltration of certain immune types. Overall, it is an exceptional experimental approach to understand responses and resistance to immunotherapies as the molecular mechanisms driving or inhibiting effective anti-tumor response can now be examined withing the cellular context.

In this study, we used ST to investigate the tumor related features that explain response or resistance to CABO/NIVO. The ST profiling of responder and non-responder samples identified transcriptional signatures that are associated with immune activation and metabolism, respectively. The intercellular interaction analysis shows that in responders these cancer-immune communications lead to B cell activation while in the areas of cancer-cancer associated fibroblasts (CAF) interactions there is activation of extracellular matrix (ECM) remodeling genes. The molecular and cellular analysis with ST indicates that mechanisms of response depend on the infiltration of immune cells into the HCC TME combined with the ability of cancer cells to trigger the immune response. On the other hand, non-responder samples are poor in immune cells because of unsuccessful antigen presentation. The ST analysis also identified remarkable HCC heterogeneity in one sample with a tumor cluster resembling responders’ samples and another cluster resembling a non-responder tumor and that is potentially associated with disease recurrence due to the expression of cancer stem cells (CSC) features not observed in other samples from our cohort. Our study demonstrates the power of ST analysis to understand responses to therapies in the context of TME composition and molecular features and identifies independent mechanisms of resistance and recurrence in HCC.

## METHODS

### Patients and Sample Acquisition

Clinical outcomes and detailed study methods for our clinical trial of CABO/NIVO were recently described by our group [10]. Briefly, we enrolled patients with potentially resectable HCC on a single-arm, open label, phase 1 clinical trial of neoadjuvant cabozantinib and nivolumab. Patients were enrolled at the Liver Cancer Multidisciplinary Clinic at the Johns Hopkins Sidney Kimmel Comprehensive Cancer Center in Baltimore. Key eligibility criteria included borderline resectable or locally advanced HCC, age greater than or equal to 18 years, an Eastern Cooperative Oncology Group performance status score of 0 or 1, and preserved liver function with a Child–Pugh score of A. The enrolled patient population included patients with high-risk tumor features that historically predict poor outcomes with upfront surgical resection such as multinodular disease, portal vein invasion, or large tumors. The trial was registered at ClinicalTrials.gov as NTC03299946 and the institutional review board of Johns Hopkins University approved the protocol. All enrolled patients provided written informed consent. Cabozantinib and nivolumab were provided by Exelixis and Bristol Myers Squibb, respectively.

A total of 15 patients were enrolled in the clinical trial, of whom 12 patients achieved successful R0 resection. Of these 12 patients, 5 patients achieved a major or complete pathological response (defined as 90% or greater tumor necrosis in the surgical resection specimen) and were classified as pathologic responders. The remaining 7 patients were considered non-responders [10]. Tumor samples from all surgical resections were immediately submerged in Optimal Cutting Temperature compound (OCT) and immersed in liquid nitrogen for quick freezing. All OCT frozen samples were stored at −80C until use.

### Spatial transcriptomics data generation

To prepare the ST slides, the samples preserved in OCT were sectioned, stained with hematoxylin and eosin, and examined by an experienced pathologist (R.A.). The areas for ST analysis were chosen based on tumor viability, presence of stroma with immune infiltration and cancer associated fibroblasts when possible. For each sample a 6 x 6mm region with those characteristics were selected for sectioning and mounting onto the ST slides.

The ST data was generated using the commercial platform Visium (10x Genomics). Briefly, from each surgical sample a 5μm section was placed in the designated area at the Visium slide and immediately stored at −80C until use. The sections were fixed in cold methanol for 30 minutes at −20C. The fixed samples were stained with hematoxylin and eosin (H&E) and imaged using the Nanozoomer scanner (Hamamatsu) at 40x magnification. Samples were permeabilized for 30 minutes at 37°C with the Permeabilization Enzyme provided with the Visium Spatial Gene Expression Reagent Kits (10x Genomics). Following permeabilization, reverse transcription; cDNA second strand synthesis, denaturation, and amplification; and library construction were performed according to manufacturer’s instructions. All libraries were sequenced with a depth of at least 50,000 reads per spot (minimum of ~250 millions per sample) at the NovaSeq (Illumina).

### Spatial transcriptomics data analysis

Sequencing data was processed using the Space Ranger software (10x Genomics) for demultiplexing and FASTQ conversion of barcodes and reads data, alignment of barcodes to the stained tissue image, and generation of read counts matrices. The processed sequencing data were inputs for the analyses using the Seurat (version 4.1.1) [15]. Data preprocessing with Seurat involved initial visualization of the counts onto the tissue image to discriminate technical variance from histological variance (e.g.: collagen enriched regions present lower cellularity that reflects in low counts), removal of low-quality spots, and normalization with SCTransform [16]. The data dimensionality was reduced with PCA and then clustered with Leiden [17]. Cell types to clusters assignments were performed based on the most variable features (genes) and revised by a pathologist (R.A.) to confirm with the liver histological regions.

Differential expression analysis of tumor spots from responders versus non-responders was performed using pseudo-bulking on DESeq2 [18]. Genes were considered differentially expressed when log 2-fold change was at least ±2 and p-adjusted value ≤ 0.01. The gene set enrichment analysis was performed using the genes ranked according to the log-fold change (LFC) determined by the differential expression analysis. We evaluated in each of the groups, non-responders and responders, the enriched expression of genes belonging to the MSigDB Hallmark Pathways [19].

### Transcriptional factor regulatory module activity analysis

Inference of regulatory modules between transcription factors (TF) and downstream regulated genes was performed using SCENIC (version 0.11.2) [20]. The annotation of cis-regulatory motif and genome ranking database are acquired from the cisTarget database available at https://resources.aertslab.org/cistarget/databases/homo_sapiens/hg38/refseq_r80/mc9nr/gene_based/. GRNBoost2 was used to fit putative regulatory module based on the co-expression between every expressed gene and predefined TF list. Each TF-gene with Importance Measure higher than 95 percentiles are included to the module. Modules with <20 genes are excluded for the downstream analysis. RcisTarget identifies TF binding motifs enrichments and prune non-direct binding modules. Finally, regulatory module activities in every spot of Visium data are quantified by AUCell. The dependencies cisTarget, GRNBoost2 and AUCell are wrapped into SCENIC.

### Ligand-receptor cell-cell signaling network reconstruction

We used Domino (version 0.1.1) [21] to analyze signaling networks based upon gene regulatory module activities in the Visium spots. Prior to calculating signaling relationship, we subset clusters to define spatial regions for analysis. Genes that expressed less than 2.5% of spots are excluded from our analysis. Similar to the implementation of Domino for single-cell RNA-seq, we adapted this approach to spatial transcriptomics computing the spot-based Pearson correlation between regulatory modules and normalized, z-score scaled expression of receptors obtained from CellphoneDB (version 2.0) [22]. Correlations are set to zero if the receptor is targeted by the transcription factor. TF regulatory modules are signaled by receptors with Pearson correlation coefficient larger than 0.3. Cognate ligands identified by CellphoneDB are required for all receptor to be included in the signaling network.

### TCGA and CIBERSORT analysis

Gene-level RNA sequencing (RNAseq) data was downloaded from Genomic Data Commons harmonized database for The Cancer Genome Atlas Liver Hepatocellular Carcinoma (TCGA-LIHC) using the TCGAbiolinks package (v2.26.0) [23]. As gene expression, we used gene counts from STAR alignment that were log2-tranformed for the further analysis. We also used CIBERSORT predicted cell proportions for TCGA-LIHC (n=371) from Thorsson et al. [24]. We used Spearman correlation to find association between gene expression and T cell proportions estimated by CIBERSORT. The analysis was done using R/Bioconductor computational environment (v4.2.2). The heatmap was plotted using ComplexHeatmap (v2.14.0) [25]

### Data and code availability

The ST data generated will be deposited on dbGAP (raw data) and GEO (processed data) and the code will be available on GitHub.

## RESULTS

### Spatial transcriptomics identifies HCC cell composition differences between responders and nonresponders to immunotherapy

We profiled HCC samples obtained from patients on a recently reported clinical trial of neoadjuvant CABO/NIVO using ST to examine tumor intrinsic molecular and cellular features of response and resistance to the treatment. Among the 12 resection specimens analyzed, 5 had a major or complete pathologic response. ST was performed for all 12 frozen surgical HCC specimens (**Figure 1A**), of which 7 (4 responders and 3 nonresponders) passed pre-determined quality control parameters (**Figure 1B** and **1C**). For 5 samples, due to extensive necrosis of the tumor, the sequencing data did not pass the quality control (low number of counts and genes detected per ST spot) for the analysis. Unsupervised clustering of the gene expression data obtained from the ST approach recapitulates the sample architecture with gene expression clusters mapping to their respective major cell types and cell subtypes commonly present in HCC samples: cancer cells, cancer associated fibroblasts (CAFs) and immune cells (**Figure 1B** and **1C**). The marker genes identified during the clustering analysis are established markers for these cell types (**Supplemental Figures 1** to **7**). The cell types assigned to the spatial gene expression clusters were confirmed by a pathologist (R.A.).

**FIGURE 1.**
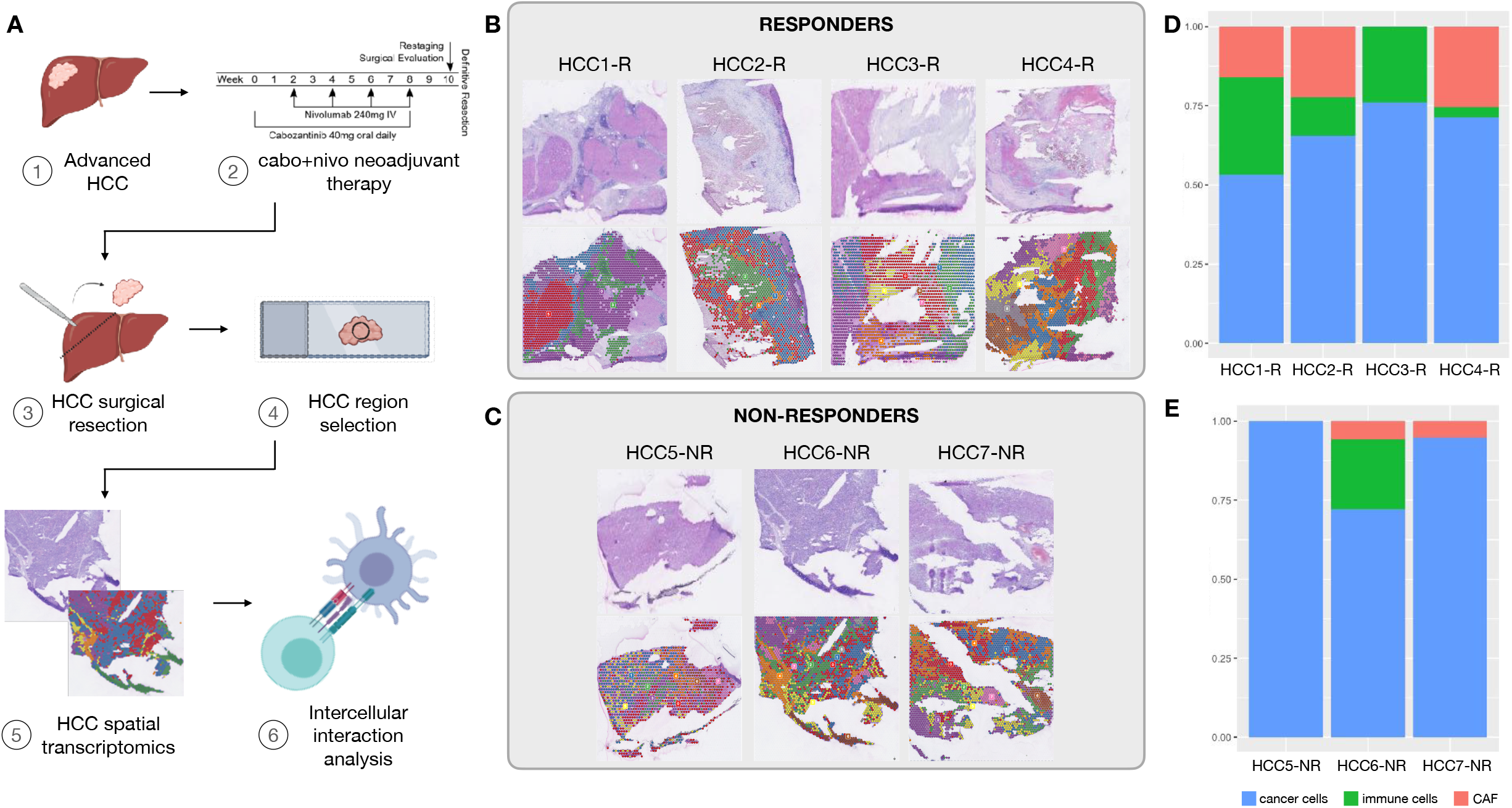
Spatial transcriptomics analysis of HCC samples treated with cabozantinib and nivolumab. A. Experimental workflow. B. Hematoxylin and eosin (H&E) stained images of the samples profiled and the spatial clusters for responders. C. H&E and spatial clusters for non-responders. D. Tumor, immune and cancer associated fibroblasts composition in each responder samples as determined by spatial transcriptomics. E. Tumor, immune and cancer associated fibroblasts proportions in non-responders samples.

Moreover, the cell type composition between responders and non-responders differs, with an increased presence of immune cells in the first group while the latest present with higher abundance of cancer cells (**Figure 1D** and **1E**). This observation corroborates the previous spatial and single-cell proteomics profiling of these samples that showed enrichment for cancer cells in non-responders and higher infiltration by immune cells in responders [10, 26]. In addition, we were able to map the CAF clusters in all 4 responders and one non-responder. The prevalence of HCC cells among non-responders suggests that the lack of response to neoadjuvant CABO/NIVO therapy is tightly correlated with the absence of immune cells infiltrating the tumor.

### Gene expression analysis of tumor compartments shows activation of immune related pathways in responders and of cell growth pathways in non-responders

To investigate the tumor intrinsic molecular changes associated with response and resistance to the neoadjuvant treatment, we performed differential expression analysis on the ST clusters mapping to HCC cells only. As the ST approach provides genome wide information, it is a suitable technology to discover the distinct transcriptional signatures between cancer cells from responders and non-responders. Additionally, the ability to map these signatures to the tissue architecture allows the selection of the ST spots that map to HCC cells for a controlled analysis of the cancer components in each sample.

The clusters mapping to tumor areas (**Figure 2A** and **2B**) were extracted from the ST data and pseudo-bulked for the differential expression analysis between responders versus non-responders. A total of 508 genes are up-regulated (**Figure 2B**, blue dots) in responders and 47 genes are up-regulated in non-responders (**Figure 2B**, green dots). The top differentially expressed genes in responders are immune related genes (e.g.: *CCL19*, *CXCL14*, *IGHM*, *CXCL6*, etc), while in the non-responders there is increased expression of tumor markers (e.g.: *AFP*, *IGF2*, *WNK4*, etc), suggesting that in patients responding to CABO/NIVO there is an active immune or inflammatory response not observed in samples from non-responders.

**FIGURE 2.**
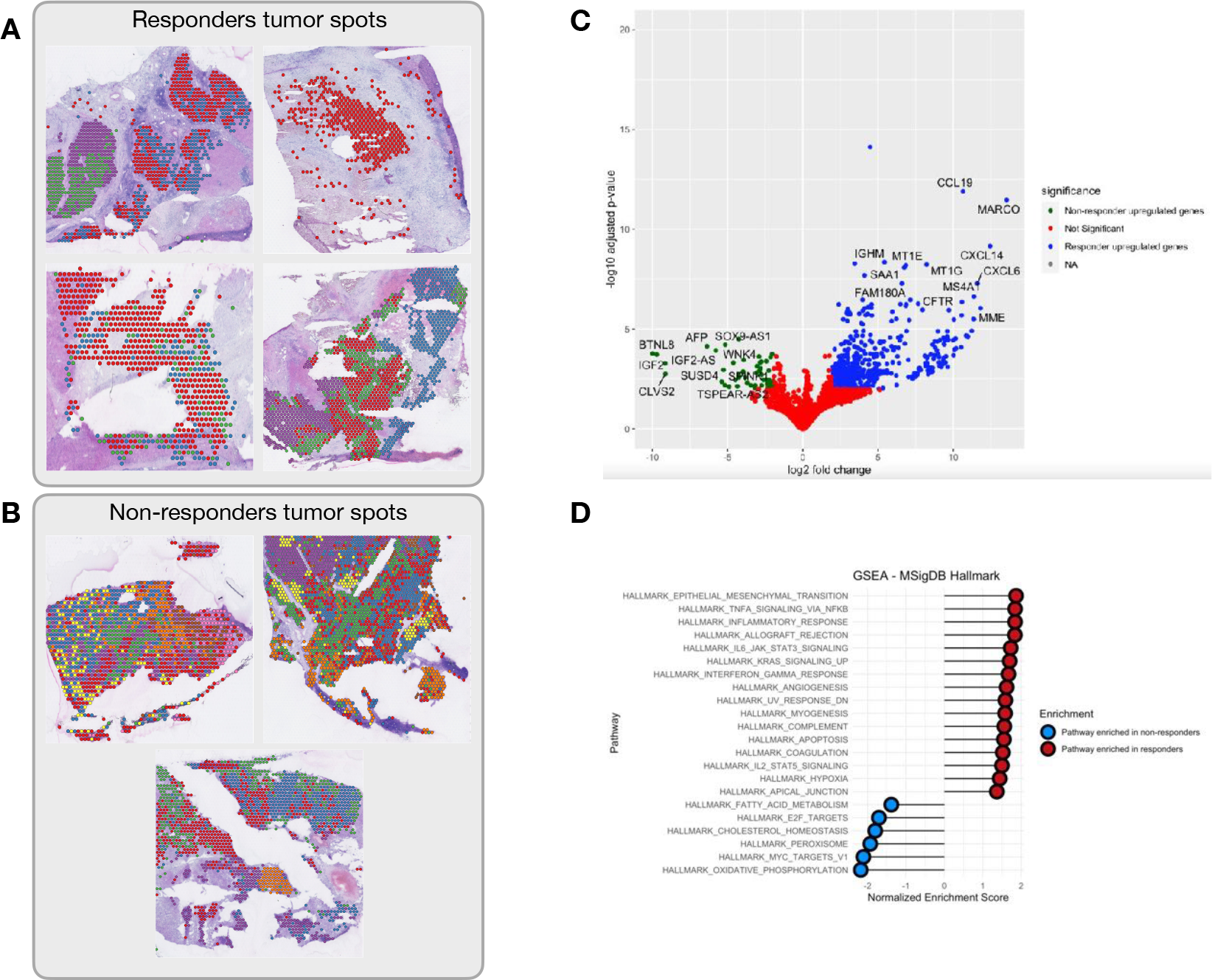
Differential expression analysis of (A) tumor clusters from responders versus (B) tumor clusters from non-responders across all patients. C. Volcano plot of the differential expression analysis showing the most differentially expressed genes in responders (blue dots) and non-responders (green dots), showing up-regulation of immune genes in responders relative to the up-regulation of hepatocellular markers among non-responders. D. Pathway enrichment analysis between responders (red dots) and non-responders (blue dots) reveal activation of immune related pathways in responders tumors, while non-responders tumors have activation of proliferation and metabolic pathways.

Subsequently, gene set enrichment analysis was performed to identify pathways enriched in responders HCC samples versus non-responders. Samples from patients that responded to CABO/NIVO are enriched for the expression of genes that belong to pathways associated with active immune response (HALLMARK_INFLAMMATORY_RESPONSE, HALLMARK_TNFA_SIGNALING_VIA_NFKB, HALLMARK_ALLOGRAFT_REJECTION, HALLMARK_COMPLEMENT) (**Figure 2C**), suggesting that among responders the tumor intrinsic features can trigger an immune response. On the other hand, non-responders lack the expression of the immune related pathways and are transcriptionally enriched by genes from signaling pathways that are involved in maintaining cell proliferation (HALLMARK_E2F_TARGETS, HALLMARK_MYC_TARGETS_V1) and metabolism (HALLMARK_OXIDATIVE_PHOSPHORYLATION, HALLMARK_CHOLESTEROL_HOMEOSTASIS), suggesting that non-responders cancer cells have their growth features maintained and activated (**Figure 2D**).

Overall, the gene expression analysis suggests that in HCC samples that did not respond to CABO/NIVO the cancer cells are not affected by the treatment and are able to maintain their proliferative and metabolic functions. On the other hand, the samples from patients that responded to the therapy show that in their cancer cells these functions are overcome by the activation of immune related pathways that could explain the tumor immune infiltration and tumor shrinkage in response to the ICI component of the treatment.

### Intercellular interaction analyses identify transcription factor regulatory networks associated with response and resistance to CABO/NIVO

Intercellular interaction analyses are facilitated for ST datasets since tissue architecture is maintained simplifying the selection of neighboring cell types to examine the potential cell to cell communication. The active interaction between neighboring cell types is critical to understand cancer biology and response to therapies since the TME has great influence on tumor biology, progression and response to therapies. Examining these cellular relationships and the molecular outcomes is critical to determine potential pathways of intervention for alternative therapeutic options with increased efficacy. The interaction analyses were performed to understand the cell-to-cell crosstalk of cancer-immune and cancer-CAF cells in response to the CABO/NIVO neoadjuvant treatment. For each of the samples, we used the identified spatial clusters and used the Domino software to determine the signaling pathways that are activated because of the intercellular interactions. Due to HCC intrinsic high heterogeneity, interaction analysis was performed for each patient individually.

Among the responders, we observed the activation of the PAX5 Domino module in the immune regions adjacent to tumor clusters (**Figure 3A**), while FOS and JUN modules are highly active in CAFs surrounding the tumor spots (**Figure 3B** and **3C**). PAX5 is a transcription factor that is central to B cells differentiation [27, 28]. The PAX5 activity co-localizes with spots showing high expression of *CD19*, *CD22*, *CD79A* and *CIITA* (B cell markers) (**Figure 3D**) confirming that this is an essential factor for B cell lineage activation and maturation. Moreover, it suggests that B cells are a critical component of the tumor immune response in HCC activated by the CABO/NIVO neoadjuvant therapy. The activation of FOS and JUN from the cancer-CAF interactions is related with high expression of ECM remodeling markers (*COL1A1*, *COL3A1*, *VIM*) (**Figure 3E**).

**FIGURE 3.**
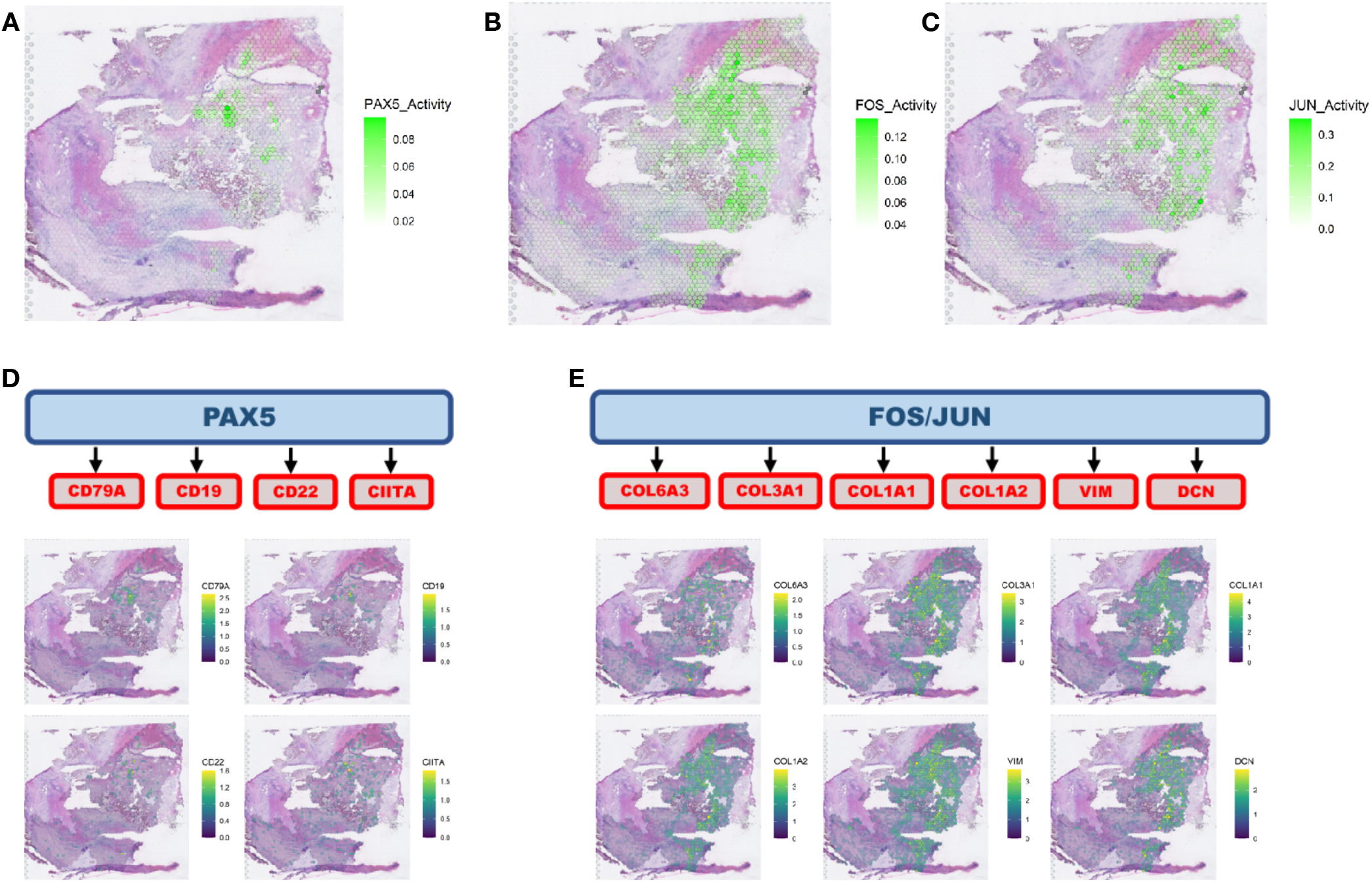
Intercellular interaction analysis. A. Cancer-immune interaction analysis identified the activation of PAX5 network. B. and C. Cancer-CAF interaction analysis pointed to the activation of FOS and JUN networks. D. PAX5 is a transcription factor that regulate B cell activity and the PAX5 network identified co-localizes with the distribution of B cells as determined by the spatial distribution of B cell markers. E. FOS and JUN are transcription factors that can regulate genes involved in extracellular matrix remodeling and the networks regulated by these genes colocalize with CAF marker genes.

This spatial distribution of tumor immune response and ECM remodeling networks suggest that in the presence of immune cells there is an active response mostly driven by B cells that could be the initial trigger for tumor cell killing by cytotoxic immune cells and recruitment of other effector immune cells. The presence of the stroma and active remodeling with increased collagen production (*COL1A1*, COL3A1) could be a result of fibrosis as a response to cell death that recruit CAFs and initiate immune exclusion in collagen/CAF rich regions and so creates a niche characterized by lack of immune cells. These findings suggest that drugs that initiate or maintain B cells activity combined with CAF inhibitors could be alternatives to increase efficacy of immunotherapies to treat HCC.

### Spatial transcriptomics analysis reveals specific HCC heterogeneity related with recurrence after neoadjuvant therapy

Among the 5 patients that responded to CABO/NIVO neoadjuvant therapy, four patients remain without disease recurrence at least 3 years after surgery, whereas a single patient developed recurrent disease after one year of therapy. We utilized ST to investigate distinct features of this single clinical outlier (hereby named HCC1-R) that might explain the unexpected early recurrence observed in this patient. The initial pathology examination of this sample identified striking histology heterogeneity, unlike any other sample in the cohort. Two distinct tumor regions were apparent in this sample: one that is immune rich (**Figure 4A**, cyan) and another that is immune poor (**Figure 4A**, dark blue). The histological distinction is recapitulated by the spatially resolved gene expression profile that identify these two tumor regions by distinct ST clusters (**Figure 4B**). As highlighted previously, the major advantage of ST is that it provides genome-wide gene expression while maintaining tissue architecture, thus an ideal approach to analyze such a unique sample.

**FIGURE 4.**
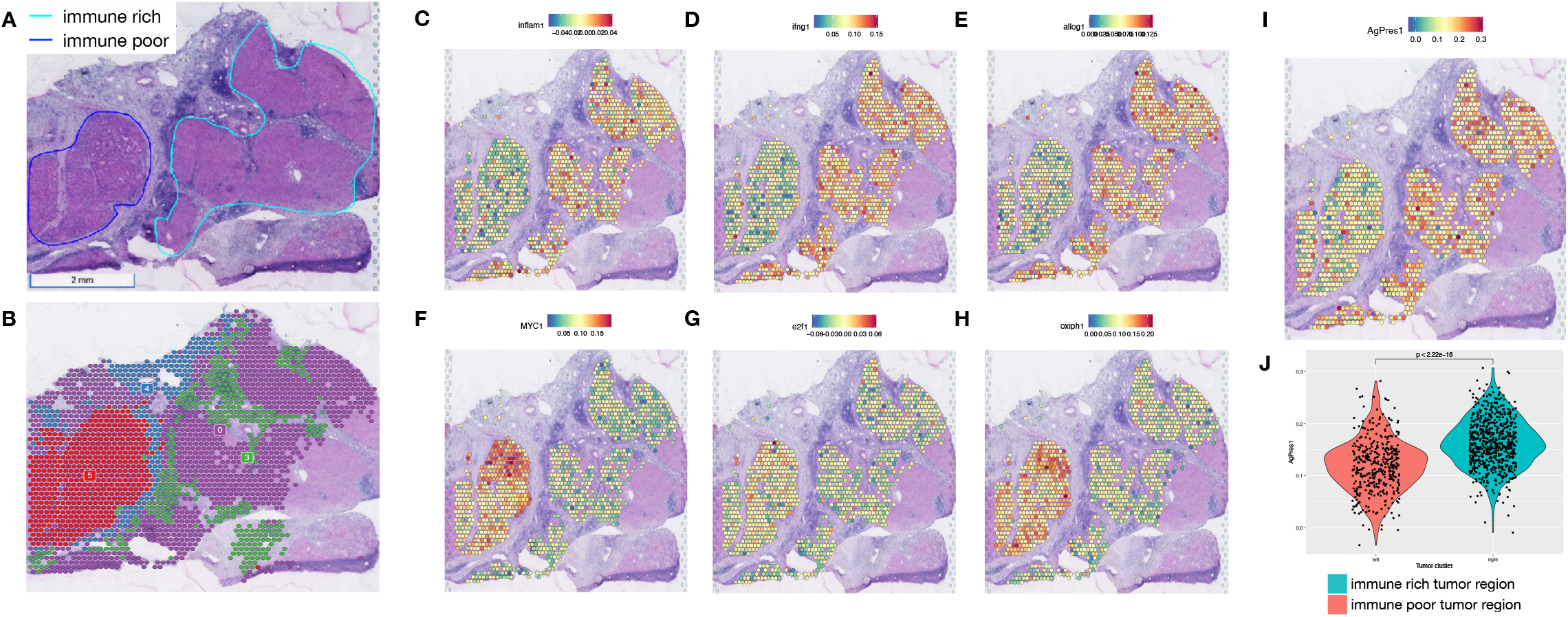
Intra-sample heterogeneity analysis with spatial transcriptomics. A. Responder sample with remarkable heterogeneity with two tumor regions, one immune-rich (cyan) and one immune-poor (dark blue). B. Each of these regions present distinct transcriptional profiles as determined by clustering analysis. C, D and E. The immune rich tumor region highly expresses immune related pathways that were initially observed to be enriched across responders’ samples. F, G and H. Proliferation and metabolic pathways, that are enriched across non-responders’ tumors, are expressed in high levels at the immune poor region. I. The immune rich tumor region expresses high levels of the antigen processing and presentation machinery genes, which is associated with more efficient attraction of immune cells.

We performed differential expression analysis to compare the immune rich and poor regions to identify differences that could explain the distinct immune infiltration between these two tumor regions. The analysis shows that among the top up-regulated genes from the immune rich region, the majority are immune markers while the immune poor region expresses increased levels of HCC specific tumor markers (**Supplemental Figure 8**). The gene set enrichment analysis, similar to the analyses of all responders versus non-responders tumors, reveals the enrichment for immune related pathways in the tumor region that is highly immune infiltrated and for growth related pathways in the tumor area that is not infiltrated by immune cells (**Supplemental Figure 8**). The expression and pathway analysis suggest that the immune rich region from the sample collected from patient HCC1-R recapitulates the transcriptional profile observed across all responders tumors while the immune poor area is similar to non-responders cancer cells.

To test our hypothesis that the two distinct regions resemble a responder (immune rich) and a non-responder (immune poor) samples, we examined if the expression of pathways enriched across responders and non-responders is reproducible on the immune rich and immune poor regions, respectively, of patient HCC1-R. Using a module score analysis of the signaling pathways enriched in both patient groups (responders and nonresponders), we mapped the averaged expression of the set of genes belonging to each of the enriched pathways. Thus, we determined if a pathway is enriched in a spot if the module score is high. The significantly enriched MSigDB pathways in responders are highly expressed in the immune rich tumor region of patient HCC1-R (**Figure 4C**, **4D** and **4E** and **Supplemental Figure 9A, B, C, D** and **E**), while those enriched in patients from non-responders are up-regulated in the immune poor tumor region (**Figure 4F**, **4G** and **4H** and **Supplemental Figure 9F, G, H, I** and **J**), thus, suggesting that the HCC heterogeneity in this sample recapitulates the features of tumors from responders (immune rich) and non-responders (immune poor). To verify if immune infiltration was associated with more efficient antigen processing and presentation, we also examined the module score for the expression of the antigen processing and presentation machinery (KEGG_ANTIGEN_PROCESSING_AND_PRESENTATION). At the immune rich tumor region the module score for this pathway is significantly higher when compared to the module score at the immune poor region (**Figure 4I**), suggesting that one of the mechanisms driving the immune cell infiltration is effective antigen processing and presentation. Finally, the intercellular interaction analysis recapitulates what was observed in other responders. PAX5 module is more active in the immune rich region, whereas FOS and JUN modules have significantly higher activation in CAFs (immune poor region) (**Figure 5A and B**). The activation of PAX5 is again associated with the areas enriched for B cells but the ECM activity is not frequent at those regions (**Figure 5C**).

**FIGURE 5.**
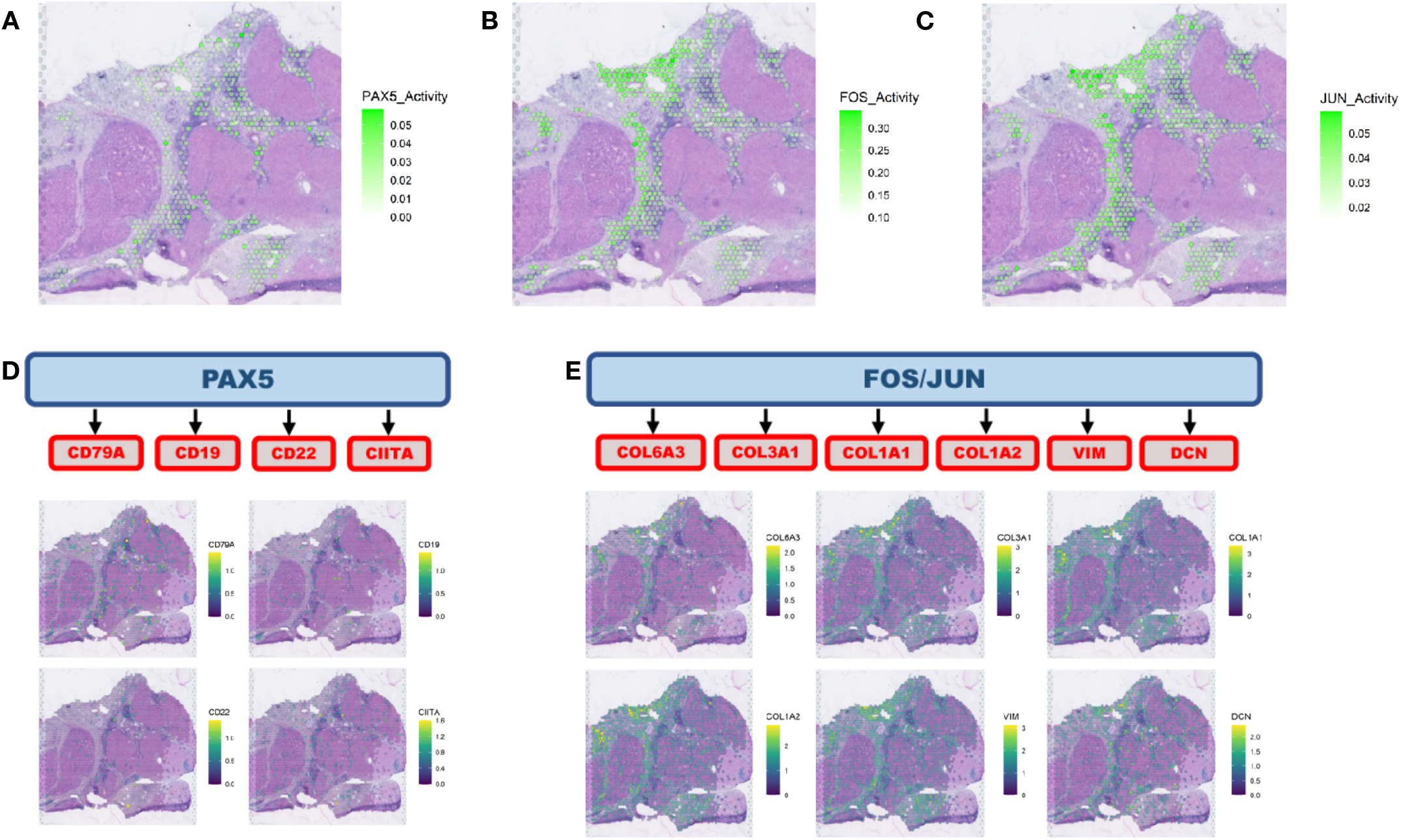
Intercellular interaction analysis in the context of intra-sample heterogeneity. A. Intercellular interaction analysis in immune rich tumor region (cancer-immune) reveals activation of PAX5. B and C. FOS and JUN are the active networks from the interaction analysis at the immune poor tumor region (cancer-CAF). D. PAX5 activation co-localizes with the expression of B cell markers concentrated at the borders of the immune rich tumor region. E. FOS and JUN networks are co-expressed with CAF markers adjacent to the immune poor tumor region.

To understand the lack of immune infiltration into one of the tumor regions from patient HCC1-R, we looked at signatures associated with immune evasion. Interestingly, a signature associated with immune evasion and resistance to different types of cancer therapies, including immunotherapies, and that is highly expressed by the immune poor tumor region is from cancer stem cells (CSC) (**Figure 6A**) [29]. CSCs are associated with therapeutic resistance, immune escape, recurrence and metastasis. The presence of HCC cells with stemness features have been previously observed in HCC and also associated with therapeutic resistance [29].

**FIGURE 6.**
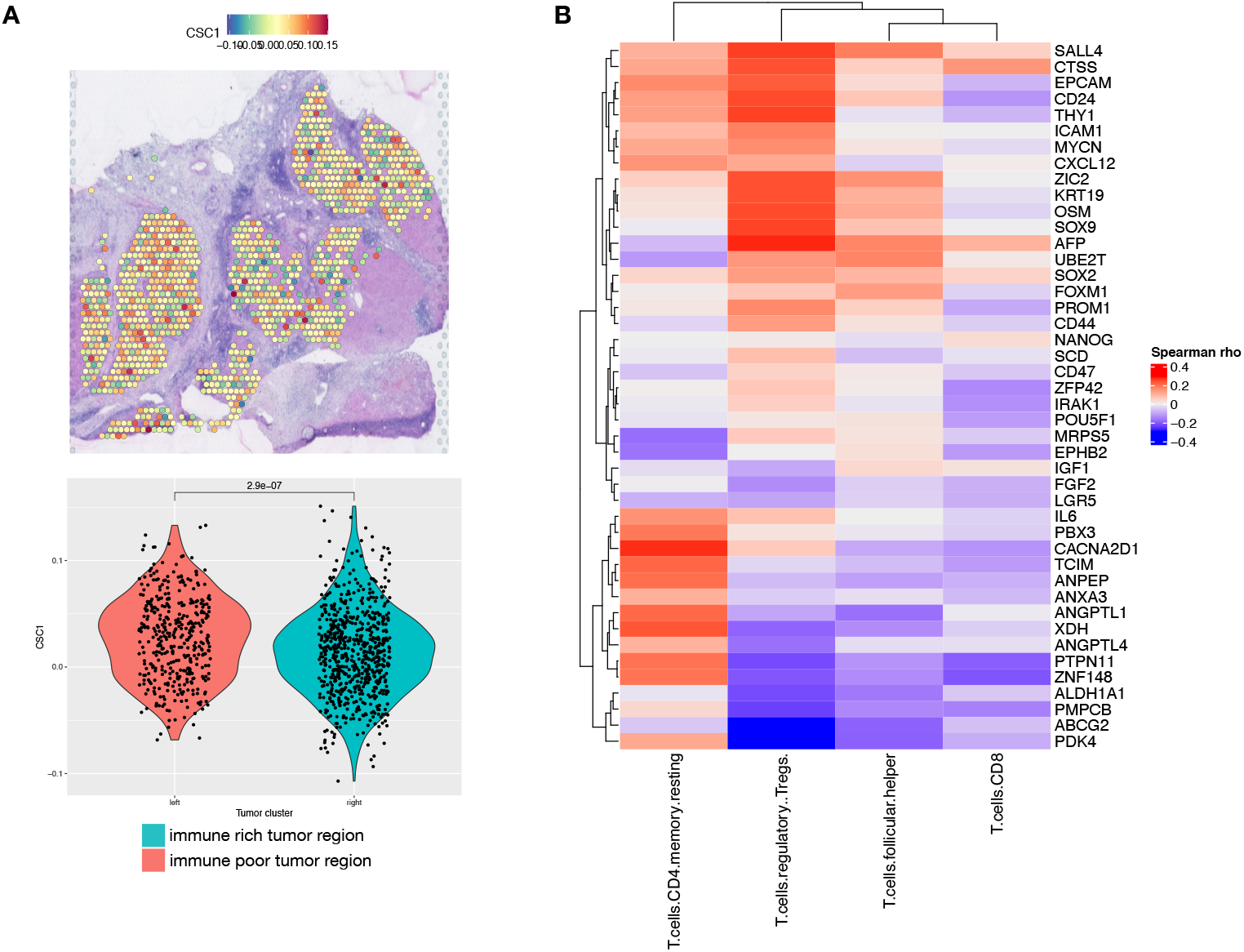
Cancer stem cells detection in hepatocellular carcinoma heterogeneous sample. A. Cancer stem cell markers are highly expressed at the immune poor tumor region of the responder sample with intrasample heterogeneity as noted by the spatial expression and the levels of expression are significantly different between the distinct tumor regions. B. Hepatocellular carcinoma samples from TCGA that expresses cancer stem cell markers are highly infiltrated by T CD4+ regulatory cells while T CD8+ cells are rare.

To validate that the CSC molecular signature is associated with HCC that fails to elicit an effective immune response and so is more prone to resistance to ICIs, we identified HCC samples from the Cancer Genome Atlas (TCGA) that express the CSC signature. Using CIBERSORT we determined the immune cell proportions in samples with high versus low expression of CSC genes. The HCC samples that expressed the CSC markers were highly infiltrated by T regulatory cells (Tregs), an immunosuppressive cell type. Samples with low expression of the CSC markers were more infiltrated by T CD8^+^ cells, that are associated with better outcomes and prolonged response to ICIs (**Figure 6B**). This difference in immune cell types that are infiltrated in CSC rich HCC relative to HCC without CSC features corroborates the hypothesis that CSCs are able to evade the immune system. The presence of CSC in the only patient that recurred from the CABO/NIVO adjuvant therapy suggests that the presence of these cells is associated with therapeutic resistance and that this population will then persist and lead to recurrence.

## DISCUSSION

Modern systemic treatment for HCC combining anti-VEGF therapy with ICI therapy prolongs survival and is widely used in the first line treatment of patients with HCC [5], however little is known about mechanisms of response and resistance to this new treatment paradigm. Prior analysis from our group using samples from a clinical trial of neoadjuvant CABO/NIVO demonstrates that clinical responses are associated with higher infiltration of immune cells while resistance was associated with diminished presence of immune cells that were in its majority immunosuppressive [10, 26]. However, due to the limited marker panels in the experimental approaches in that previous study, the molecular pathways that underlie immune response and changes in the cancer cells themselves were not investigated to determine their role in the response to the neoadjuvant therapy. ST analysis is a novel approach that enables whole transcriptomics analysis within the tissue architecture context. We applied ST to unveil the cancer cells intrinsic features that are associated with response and resistance to CABO/NIVO. Since the ST approach used to profile the specimens collected is genome-wide, the analysis is not restricted to markers or pathways previously selected.

Using ST, we identified gene expression clusters that map to specific cell types in the HCC samples. With that, gene expression analysis of the subset of cancer cells from each patient demonstrate that in responders there is an active anti-tumor immune response characterized by the high expression of immune related genes and recruitment of immune related pathways. Moreover, in non-responders cancer cells we observe the activation of proliferation and metabolic pathways. The intercellular interaction analysis of cancer-immune cells demonstrates that the immune response observed is driven by a strong B cell response. The cancer-immune cell interaction is associated with activation of a PAX5 module that is enriched in regions with increased presence of B cells. PAX5 is a transcription factor essential for B cells maturation, commitment, immunoglobulin rearrangements and activity [27]. In tumors, B cells are frequently found in tertiary lymphoid structures where they act by presenting antigens to T cells that could lead to their activation into effector cytotoxic cells [30]. Thus, the presence of B cell activation suggests that there is also induction of a T cell cytotoxic response and effective response to treatment with an ICI, in this case nivolumab. Another observation from the intercellular interaction analysis is that in the regions with cancer-CAF communication there is activation of ECM remodeling. This was also an observation across responders samples. The active ECM remodeling in the presence of therapeutic response, indicates that in the areas of tumor cell death there is a fibrosis process that replaces the space once occupied by the cancer cells. Further analysis with temporal model or sample collection would reveal if that process will lead to future resistance to the therapy, as the presence of desmoplasia with a dense collagen ECM may create a barrier to immune cells infiltration into the remaining tumor. Overall, these findings suggest that therapeutic resistance is associated with inability of cancer cells to trigger an anti-tumor immune response that would increase immune cells infiltration and that this response rely on the presence of B cells that will prime or activate cytotoxic T cells. Thus, B cell presence is a potential cellular biomarker of response to therapies with ICIs.

With the spatially resolved analysis of this cohort of neoadjuvant treated HCC samples, it was possible to examine a sample from a patient that initially presented with therapeutic response but that later recurred. The surgical specimen collected from this patient shows a notable tumor heterogeneity pattern with an immune poor tumor region separated from an immune rich tumor region. This tissue distribution would have been lost if the samples were profiled using single-cell approaches as those depend on sample dissociation. The immune poor tumor region molecularly resembles the tumors from non-responders, while the immune rich counterpart is similar to responders’ samples that have high immune activity. The immune infiltration is a result of more effective antigen presentation by the cancer cells from the immune rich region. Successful antigen presentation is a well-known component of prosperous tumor immune response and could be used as a molecular biomarker to predict responses to ICIs [31]. On the other hand, the immune poor tumor area does not express high levels of the antigen presenting machinery and thus evades the immune system. One of the reasons that could explain the down-regulation of antigen presentation is that the cancer cells in that cluster express CSC markers. Due to their stemness features, these cells are expected to be more successful in immune escape and are also more prone to therapeutic resistance [29, 32]. Furthermore, the presence of CSC could explain the tumor recurrence in this patient since these unique cells are frequently present in disease relapse. Hence, the presence of CSC in the tumor is another biomarker that could potentially predict response to therapies and the risk of recurrence after treatment. Ultimately, with the analysis of this clinical outlier, we were also able to demonstrate a distinction between intrinsic resistance and recurrence. The CSC signature is only up-regulated in patient HCC1-R. We did not observe the CSC signature in the samples from patients that did not respond to CABO/NIVO. Thus, the mechanisms of intrinsic resistance and acquired resistance, leading to recurrence, are different.

## CONCLUSIONS

Although our study is limited to a small sample size, this is an exceptional cohort that used a therapeutic combination to treat HCC with promising clinical benefits to a fraction of patients. Regarding the experimental approach, ST also has limitations as there is no single-cell resolution due to the fact that each spot can capture the signal of more than one cell limiting the ability of specifically identifying individual immune cell types and subtypes. However, it demonstrates its usefulness since it allows heterogeneity identification in the spatial context. As previously mentioned, we would have not been able to identify the unique tumor architecture of the single responder that recurred after therapy. The analysis of that sample allowed the identification of an important tumor feature that is potentially associated with the recurrence in that specific patient. In summary, the ST analysis of HCC clinical trial samples identified molecular and cellular mechanisms of therapeutic response and resistance. The mechanisms unveiled by the spatially resolved gene expression analyses would not be found by bulk or single-cell analysis as the lack of the spatial component would not have permitted the interpretations related to the cellular distributions and interactions we were able to extract. Further analysis on other clinical cohorts are essential to corroborate our findings and could potentially identify alternative therapeutic interventions for cancer types that still do not benefit from treatment with ICIs.

## LIST OF ABBREVIATIONS

AFP: alpha fetoprotein
CABO: cabozantinib
CAF: cancer associated fibroblast
CCL19: chemokine ligand 19
CIITA: class II major histocompatibility complex transactivator
COL1A1: collagen type I alpha 1 chain
COL3A1: collagen type III alpha 1 chain
CSC: cancer stem cell
CXCL6: chemokine ligand 6
CXCL14: chemokine ligand 14
cDNA: complementary deoxyribonucleic acid
CSC: cancer stem cells
ECM: extracellular matrix
FOS: fos proto-oncogene
HCC: hepatocellular carcinoma
H&E: hematoxylin and eosin
ICI: immune checkpoint inhibitor
IGF2: insulin like growth factor 2
IGHM: immunoglobulin heavy constant mu
JUN: jun proto-oncogene
LFC: log-fold change
LIHC: liver hepatocellular carcinoma
NIVO: nivolumab
OCT: optimal cutting compound
PAX5: paired box 5
PCA: principal component analysis
PD-1: programmed cell death 1
ST: spatial transcriptomics
TCGA: The Cancer Genome Atlas
TF: transcription factor
TME: tumor microenvironment
Treg: T regulatory cell
VEGF: vascular endothelial growth factor
VIM: vimentin
WNK4: WNK lysine deficient protein kinase 4

## Ethics approval and consent to participate

All patients provided written informed consent prior to enrollment, and the trial was registered under ClinicalTrials.gov as NCT03299946. The protocol was approved by the Institutional Review Board (IRB) at Johns Hopkins University. Cabozantinib was supplied by Exelixis and nivolumab was supplied by Bristol-Myers Squibb.

## Acknowledgments

We thank the Johns Hopkins University School of Medicine Experimental and Computational Genomics Core for performing the sequencing of the spatial transcriptomics libraries and for the pre-processing of the raw sequencing data.

## Author’s Contributions

EJF and LTK instigated and supervised the study. SZ, MY, EJF and LTK planned, designed and wrote the manuscript with input from all authors. LTK and GM performed the experimental methods. SZ, LY and LD performed computational analysis and data interpretation. GM, QZ, AD, ATFB, JE, ASP and RA provided technical and material support. MY and EMJ provided clinical expertise. All authors discussed the data, contributed to the manuscript preparation, and approved the final manuscript.

## Contributors information

Shuming Zhang

Long Yuan

Ludmila Danilova

Guanglan Mo

Qingfeng Zhu

Atul Deshpande

Alexander T.F. Bell

Jennifer Elisseeff

Aleksander S. Popel

Robert A. Anders

Elizabeth M. Jaffee

Mark Yarchoan

Elana J. Fertig

Luciane T. Kagohara

