## supplemental figures for "Spatial transcriptomics analysis of neoadjuvant cabozantinib and nivolumab in advanced hepatocellular carcinoma identifies independent mechanisms of resistance and recurrence"

Supplemental Figure 1

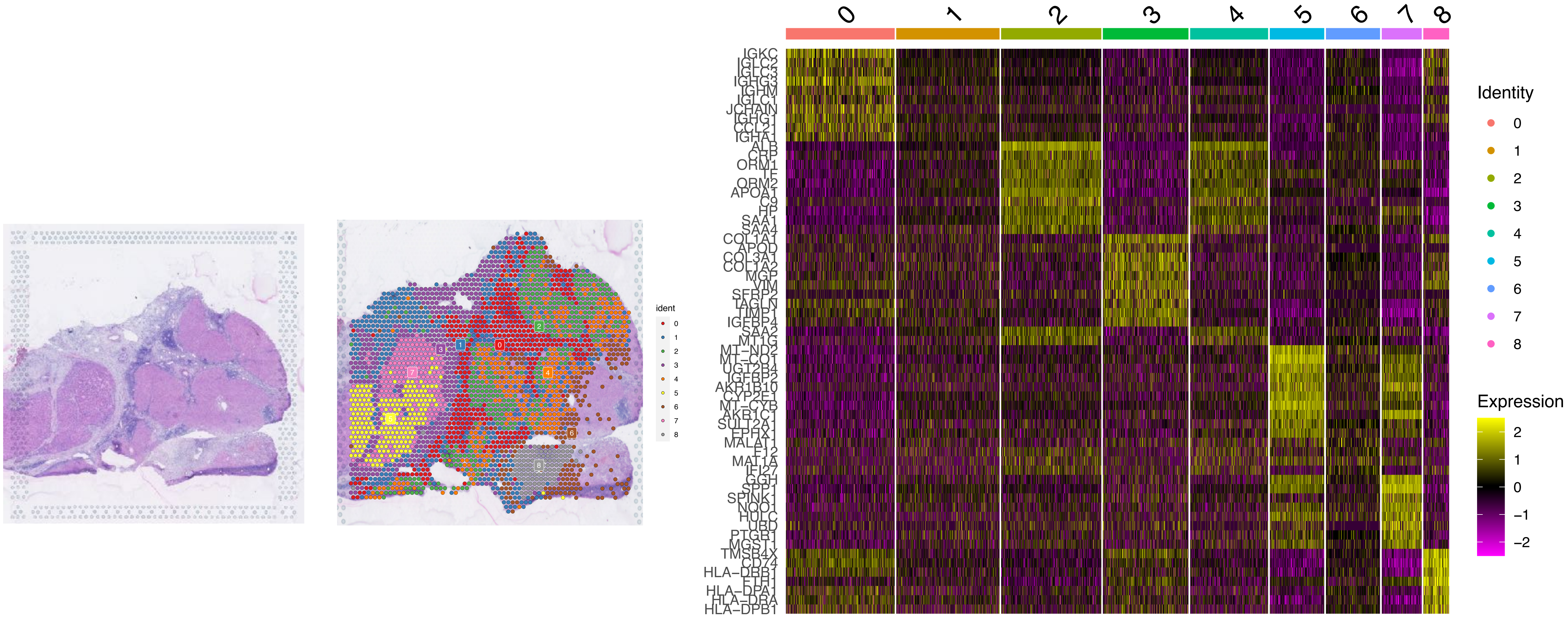


Supplemental Figure 1 - Clusters markers for patient HCC1-R. Stained spatial transcriptomics section (left panel) used for pathology examination. The unsupervised clustering recapitulates the histological composition of the the sample (middle panel). The multiple clusters are determined based on the top variable genes (right panel).

Supplemental Figure 2

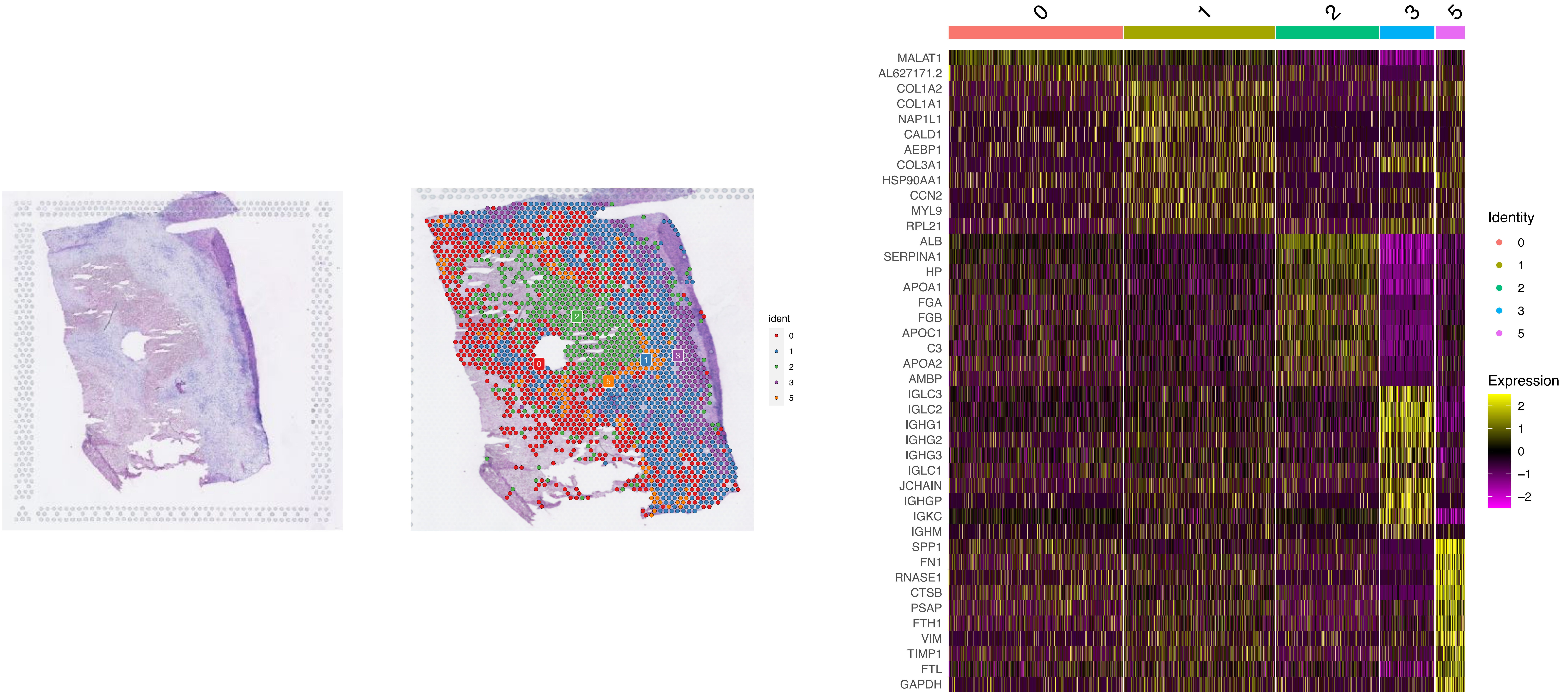


Supplemental Figure 2 - Clusters markers for patient HCC2-R. Stained spatial transcriptomics section (left panel) used for pathology examination. The unsupervised clustering recapitulates the histological composition of the the sample (middle panel). The multiple clusters are determined based on the top variable genes (right panel).

Supplemental Figure 3

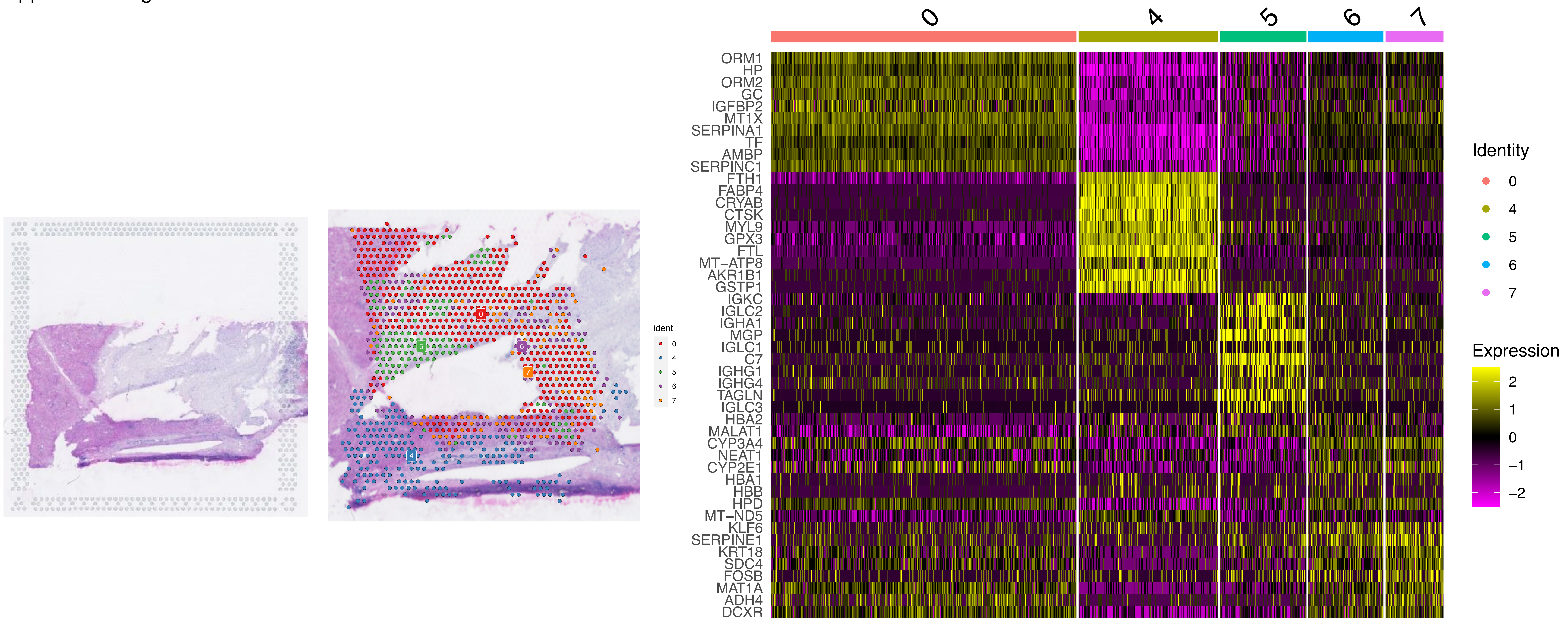


Supplemental Figure 3 - Clusters markers for patient HCC3-R. Stained spatial transcriptomics section (left panel) used for pathology examination. The unsupervised clustering recapitulates the histological composition of the the sample (middle panel). The multiple clusters are determined based on the top variable genes (right panel).

Supplemental Figure 4

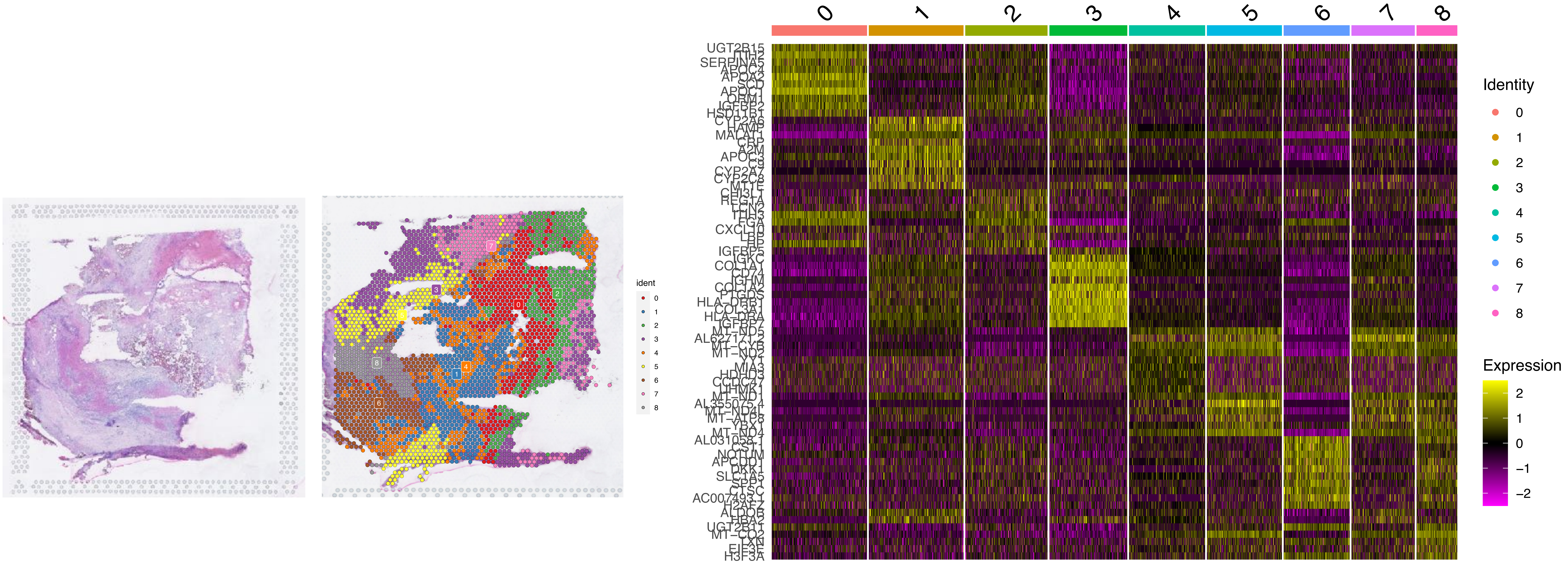


Supplemental Figure 4 - Clusters markers for patient HCC4-R. Stained spatial transcriptomics section (left panel) used for pathology examination. The unsupervised clustering recapitulates the histological composition of the the sample (middle panel). The multiple clusters are determined based on the top variable genes (right panel).

Supplemental Figure 5

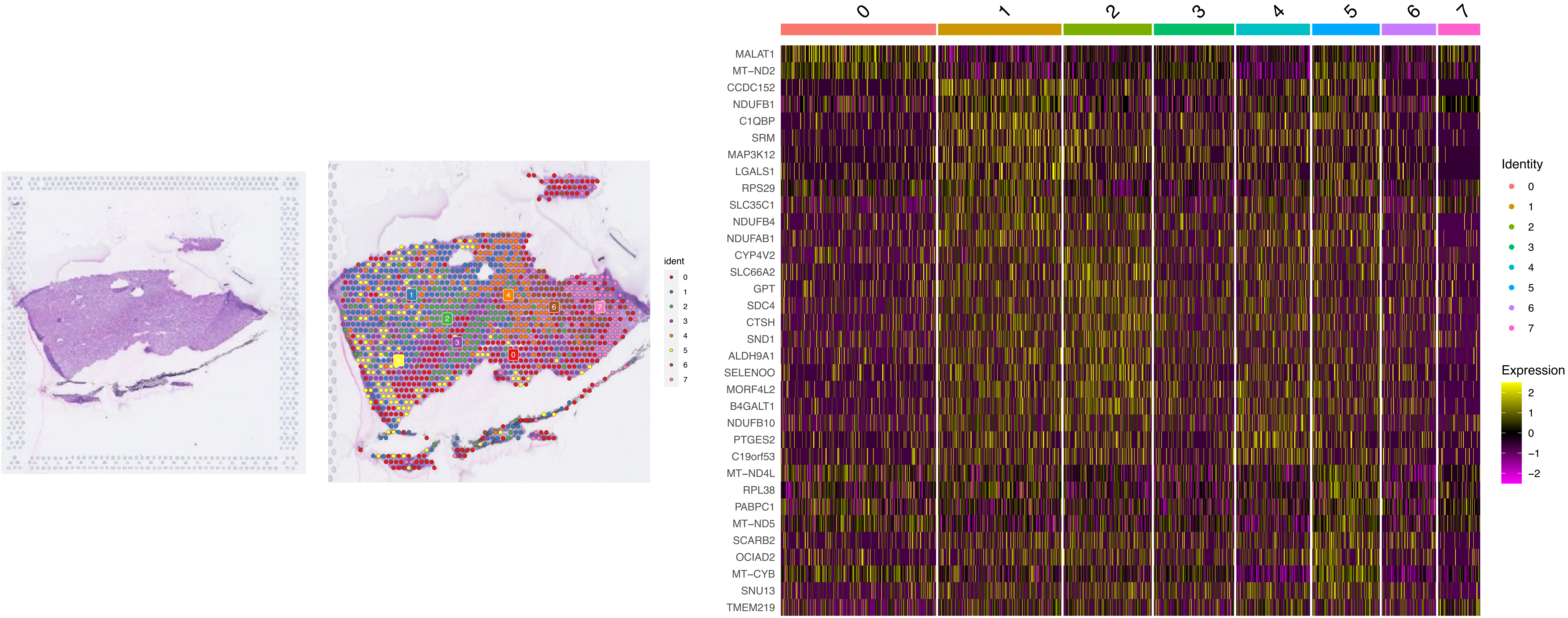


Supplemental Figure 5 - Clusters markers for patient HCC5-NR. Stained spatial transcriptomics section (left panel) used for pathology examination. The unsupervised clustering recapitulates the histological composition of the the sample (middle panel). The multiple clusters are determined based on the top variable genes (right panel).

Supplemental Figure 6

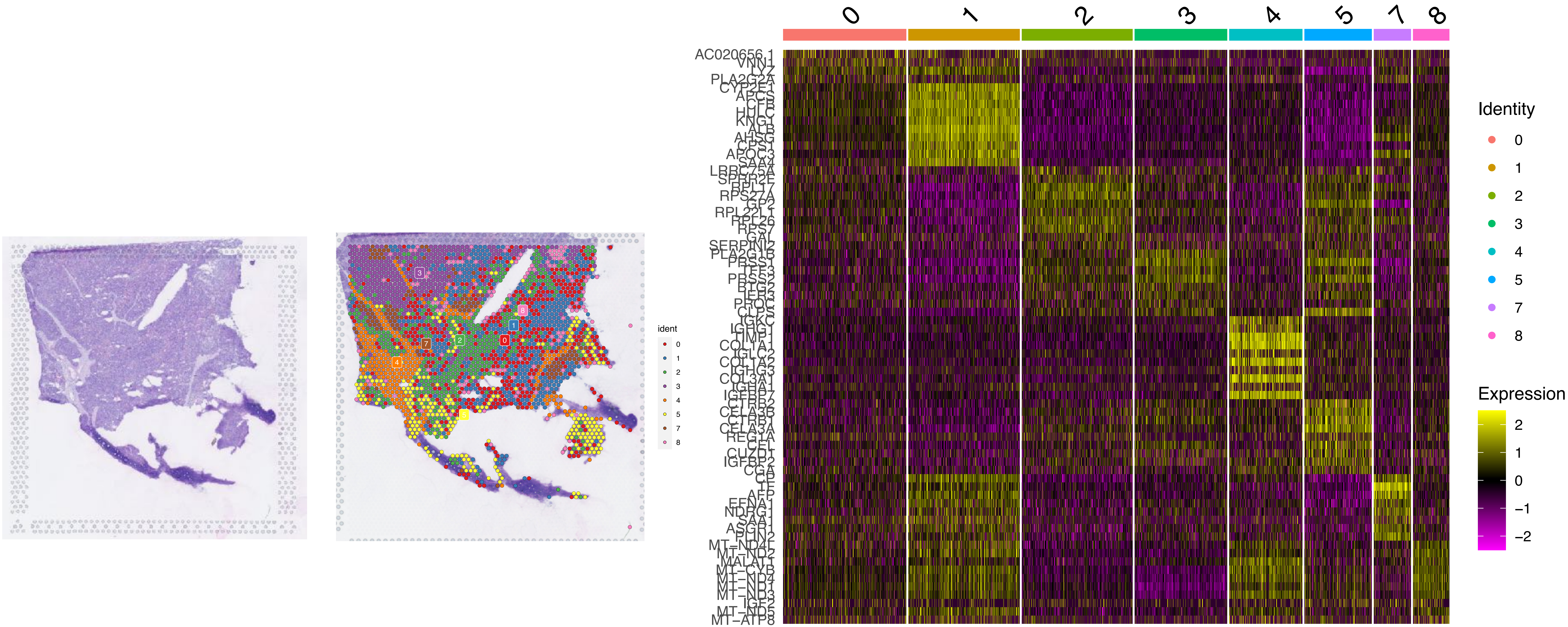


Supplemental Figure 6 - Clusters markers for patient HCC6-NR. Stained spatial transcriptomics section (left panel) used for pathology examination. The unsupervised clustering recapitulates the histological composition of the the sample (middle panel). The multiple clusters are determined based on the top variable genes (right panel).

Supplemental Figure 7

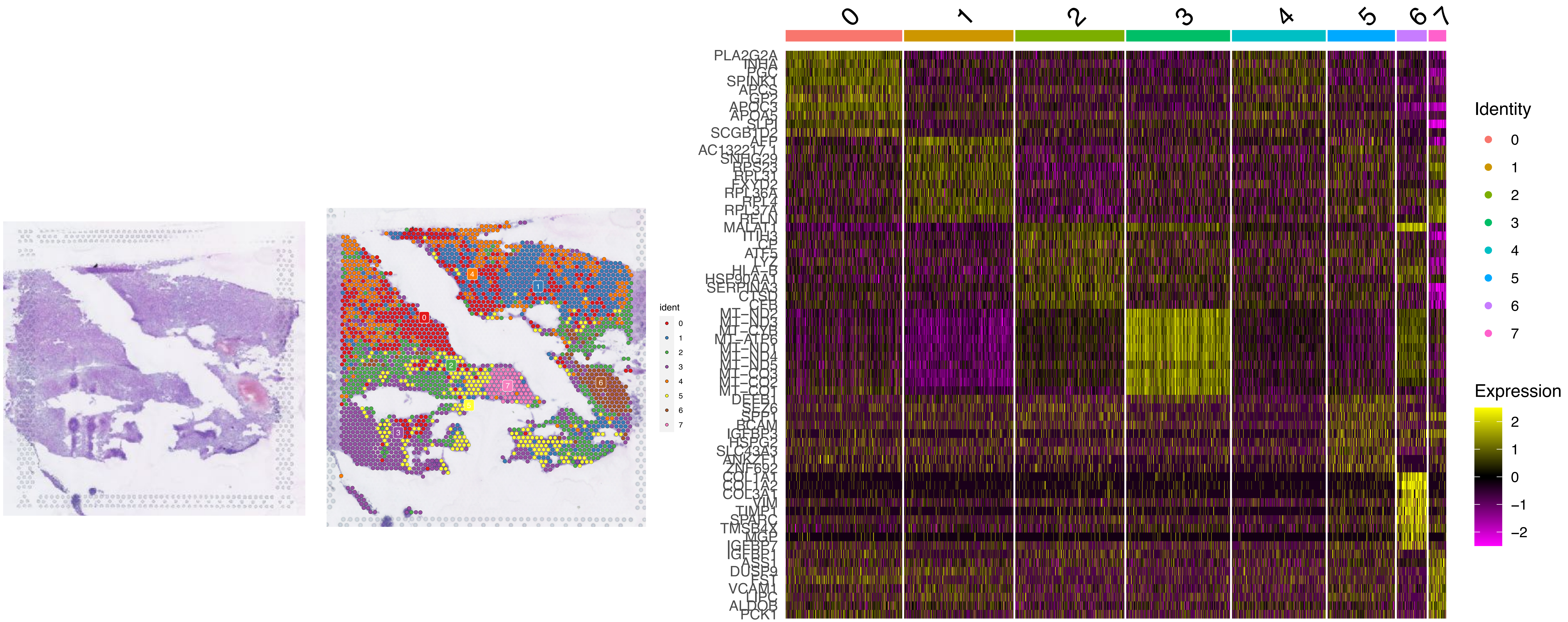


Supplemental Figure 7 - Clusters markers for patient HCC7-NR. Stained spatial transcriptomics section (left panel) used for pathology examination. The unsupervised clustering recapitulates the histological composition of the the sample (middle panel). The multiple clusters are determined based on the top variable genes (right panel).

Supplemental Figure 8

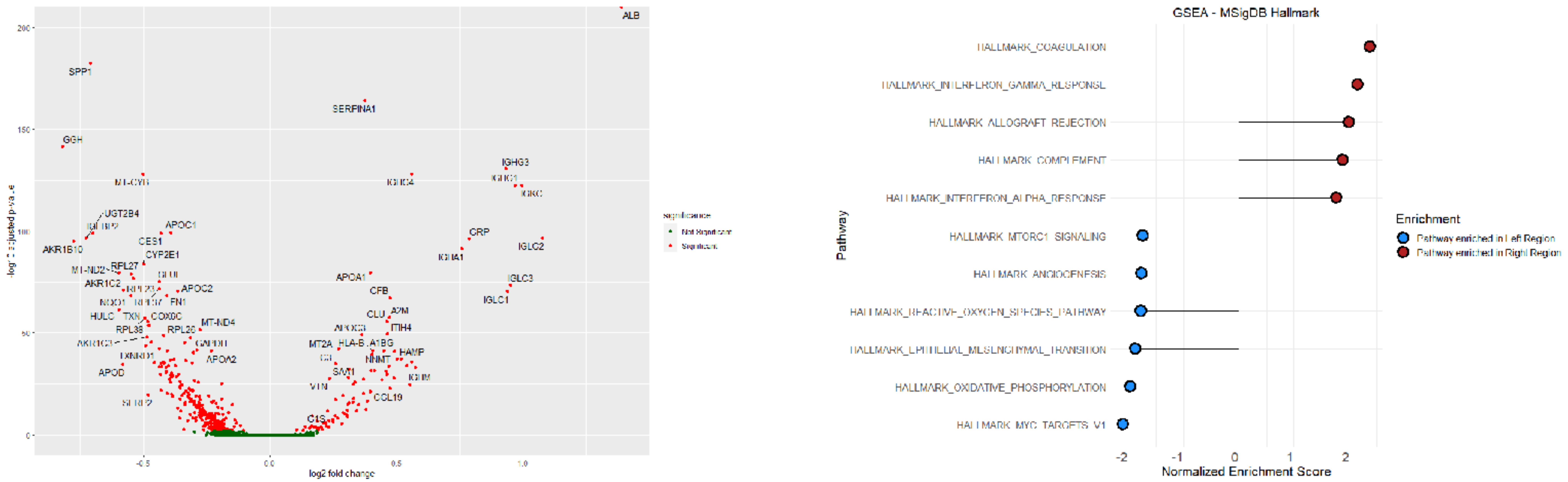


Supplemental Figure 8 - Differential expression analysis between the immune poor and immune rich regions on sample from patient HCC1-R. The volcano plot (left panel) represents the up-regulated genes in the immune poor tumor region (genes on the left) and in the immune rich tumor region (genes on the right). Most of the markers in the immune poor region are HCC or common tumor markers while the immune rich region up-regulates immune related genes. The gene set enrichment analysis (GSEA - right panel) revealed that the immune rich tumor expresses genes associated with immune response, while the immune poor tumor expresses genes from metabolic and cell proliferation pathways.

Supplemental Figure 9

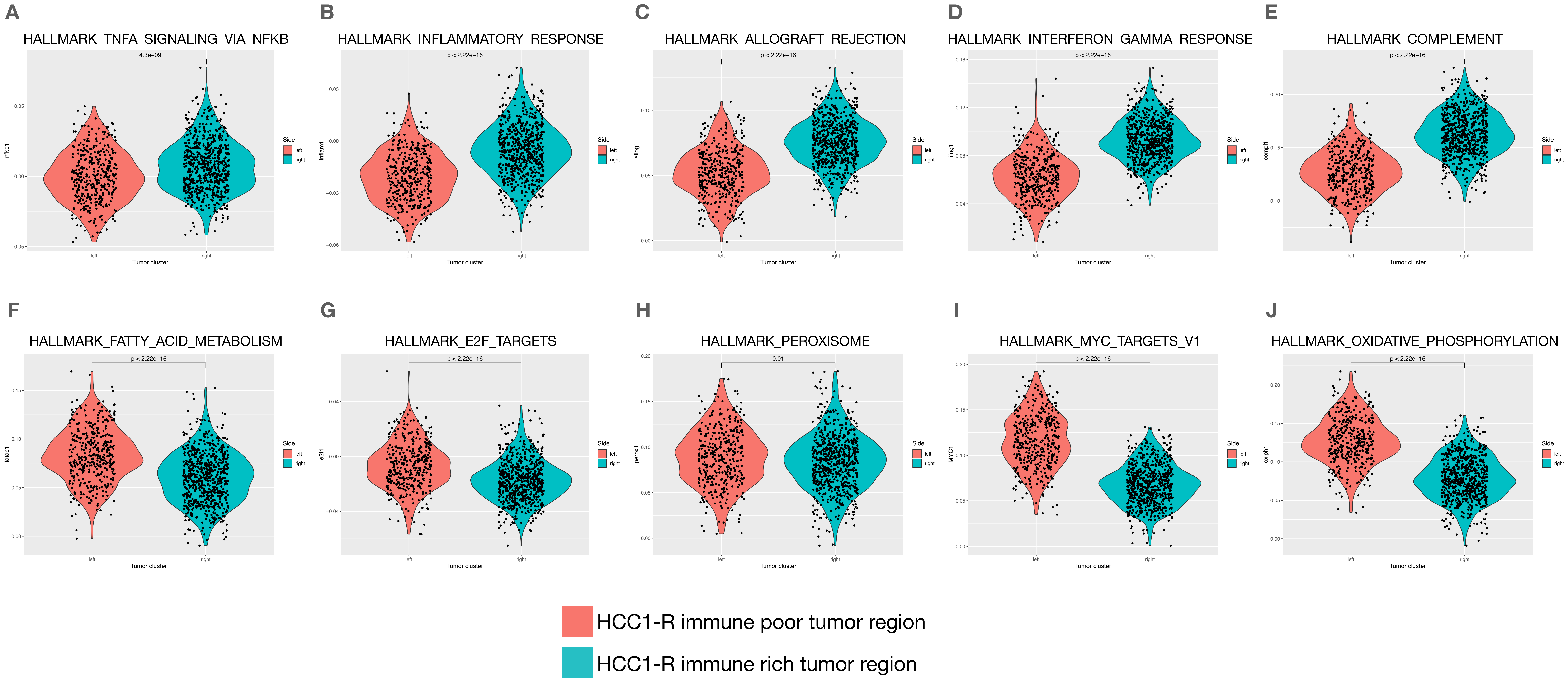


Supplemental Figure 9 - Expression of MSigDB Hallmark pathways across all spots on the immune poor and immune rich tumor regions from patient HCC1-R. A, B, C, D and E represent the module score expression distribution of immune related pathways in HCC1-R immune rich regions that are enriched in responder samples according to gene set enrichment analysis. F, G, H, I and J represent the module score expression distribution of proliferation and metabolic related pathways in HCC1-R immune poor regions that are enriched in non-responder samples according to gene set enrichment analysis.
